# Why Speech Motor Blocks Emerge in a Communicative Context: An Active Inference Model of Stuttering

**DOI:** 10.64898/2026.08.17.745328

**Authors:** Birtan Demirel, Thomas Parr, Youssuf Saleh, Eric Jackson, Timothy Denison, Sanjay Manohar

## Abstract

Adults who stutter can speak fluently when speech is not addressed to another person, but stuttering emerges when they aim to convey information to a listener. The value of the information being conveyed to the listener also affects the likelihood of stuttering. Why should the mere absence of a listener neutralise a profound motor deficit, and why does a word’s predictability affect whether it is spoken fluently?

To resolve this socio-motor paradox, we develop a computational model of stuttering within an active inference architecture. The model represents the communicative context, including whether a listener is present and whether the agent is speaking or listening. It was designed around two candidate mechanisms for stuttering, a prior for silence and rigid phoneme sequencing precision. Using both, the model produced fluent private speech and more stuttering-like events during social speech. In the same parameter regime, the model also showed more stuttering-like events on words with higher information value, and produced a word-length effect, in which disfluency increased with longer words.

To our knowledge, this is the first model of stuttering to generate both the private speech and the information-value effect from inferred communicative context. By representing the listener as a hidden state that makes the sensory consequences of resuming speech ambiguous, the model offers a computational link between social cognition and speech-motor instability, and suggests that speech fluency depends on whether the speaker believes anyone is present. Clinically, it may offer testable hypotheses and a route to personalising treatment, since the same overt severity can arise from different combinations of parameters.

## 1. Introduction

Developmental stuttering is commonly described as a disorder of speech motor control. Speech fluency can be strongly modulated by low-level sensorimotor manipulations such as altered auditory feedback (Howell, 2004), speaking in unison with another voice (Saltuklaroglu et al., 2009), metronome-paced speech (Azrin et al., 1968), singing (Glover et al., 1996), and slowed speech rate (Ingham et al., 1974). Yet one of the most striking features of stuttering is that a person who stutters can become completely fluent when speech is not addressed to another agent, without the need for sensorimotor modulation. In a carefully designed experiment, Jackson and colleagues (2021) investigated the private-speech effect, defined as speaking to oneself when alone without any communicative intent. Across more than 10,000 syllables of private speech, only seven stuttering events were measured, and the mean stuttering rate was 0.04% of syllables. By contrast, the same participants stuttered at their baseline rates when speaking in conversation, reading aloud, or producing the same utterances to listeners. This phenomenon is consistent with earlier studies reporting that stuttering is modulated by higher-level social and communicative factors, including communicative pressure, audience presence, social evaluation, and the speaker’s anticipation of how their speech will be perceived (Hahn, 1940; Perkins et al., 1991; Peters & Guitar, 1991; Quarrington, 1965). These findings challenge accounts that explain stuttering purely at the sensorimotor level, without reference to higher-level communicative processes. Consequently, a computational model must explain why the same speech motor control system becomes unstable when speech is directed to a listener.

Stuttering is also sensitive to the information value of words. Words that are less predictable from what came before carry more information, and they are more likely to be stuttered. If stuttering were only a failure of articulation, the predictability of the next word should not matter. However, cumulative evidence indicates that it does. Early studies found that less predictable words were more likely to be stuttered (Quarrington, 1965; Schlesinger et al., 1965; Soderberg, 1971; Wingate, 1979; for review, see Brundage & Bernstein Ratner, 2022). Later findings were mixed when word length, sentence position, and initial phoneme were controlled (Lanyon, 1969; Lanyon & Duprez, 1970). Warner and colleagues (2026) revisited this subject in spontaneous speech from 35 adults who stutter. Information value was quantified using GPT-2 word surprisal, while controlling for linguistic factors such as word length, sentence position, grammatical function, word frequency, neighbourhood density, and initial phoneme. Their findings showed that higher surprisal words were more likely to be stuttered, even after these covariates were included. Therefore, communicative context may act on stuttering at two levels: whether speech destabilises at all and how likely disfluency is for a given word.

Existing theoretical and computational models have made important progress on the motor control aspect of stuttering. These include asynchrony between execution and planning (EXPLAN model), which links stuttering to mistiming between two hierarchical components of the motor system (Howell & Au-Yeung, 2002), and Gradient Order Directions Into Velocities of Articulators (GODIVA), which attributes stuttering to impaired selection and initiation of successive speech-motor programs, such that the system fails to release or transition to the next syllable at the right moment (Civier et al., 2013; Guenther, 2016). This motor perspective builds on foundational work showing that fluent speech requires the timed coordination of the tongue, lips, jaw, and larynx across successive articulatory movements (Browman & Goldstein, 1992; Perrier et al., 1996; Saltzman & Munhall, 1989). More recently, behavioural and neurophysiological evidence of stuttering has drawn attention to disruptions in speech initiation. According to this view, stuttering can arise when the cortico-basal ganglia-thalamo-cortical loop fails to start articulation (Alm, 2004; Chang & Guenther, 2020; Civier et al., 2013). This view is supported by electrophysiological findings showing atypical beta oscillations around speech onset. These include stronger beta suppression during speech preparation and stronger beta synchronisation during articulation (Mersov et al., 2016), low beta activity before overt reading that correlates with stuttering severity (Korzeczek et al., 2022), and increased right pre-supplementary motor area beta power before stuttered utterances compared with fluent ones (Orpella et al., 2024). Overall, these findings suggest that stuttering involves unreliable timing in speech motor sequencing.

Despite the success of these motor models, the private-speech effect presents a fundamental puzzle for motor control research. How can a profound motor deficit vanish in the absence of a listener? We propose that resolving this paradox requires looking beyond the motor system itself. Our hypothesis is that communicative states, such as speaking or listening, are represented by a distinct variable within a predictive processing framework. Since active inference unifies sensorimotor execution, perceptual feedback, and high-level contextual beliefs under a single optimisation principle, it automatically captures the fact that an agent’s motor control is linked to their relation to the social environment (Parr et al., 2026). Active inference has recently been suggested for auditorily guided speech production and compared directly with GODIVA and state feedback control (Bradshaw et al., 2026). It has also been proposed as an account of stuttering itself, in which aberrant sensory precision inhibits syllable initiation (Usler, 2025). In our model, the presence of a listener renders the social context more uncertain, and because word-to-word transitions are conditioned on that context, the upcoming utterance becomes more ambiguous. Since action selection favours outcomes the agent can predict confidently, an agent in a particular parameter regime may withhold speech rather than initiate it.

By using this model, we first identified a parameter regime that reproduced the private-speech effect and also the higher disfluency rate of higher-surprisal words. We also reproduced the tendency for longer words to show stuttering-like behaviour more often (Howell et al., 2006; Warner et al., 2023), without any further tuning. Sweeping the two parameters that prominently define this regime mapped the surrounding landscape, showing that fluent speech occurs in a specific window.

## 2. Methods

### 2.1. Computational model

Under the active inference approach we pursue here, we assume that the brain employs a generative world model, comprising priors and likelihood distributions. This model predicts sensory data and generates action through reflexive fulfilment of proprioceptive predictions. Perceptual inference proceeds through updating priors to posteriors using standard Bayesian inference schemes. The generative model follows the partially observable Markov decision process formulation described in Parr et al. (2026), implemented as a discrete state-space active inference model solved via belief propagation. The model architecture is described in detail there; here we focus on the aspects most relevant to simulating stuttering-like behaviour with and without a conversational partner.

Briefly, the model comprises three hidden state factors that can be used to explain sensory inputs (Fig. 1). The first is a social context factor with six states, capturing whether the agent is silent and alone, silent with company, speaking to itself, speaking to another person, listening to another person, or where the agent and partner are both speaking. The second factor encodes the current word being produced, drawn from a vocabulary of 16 pseudo-words. The third factor tracks the agent’s position within the phonemic orbit of each word, which describes how far through a word the agent has progressed, from onset to completion, plus a terminal silent state that marks word boundaries. Our choice of phoneme as the primary unit aims to keep our model interpretable, and is supported by data suggesting that prefrontal neurons encode the phonetic structure of planned words before articulation (Khanna et al., 2024).

**Figure 1:**
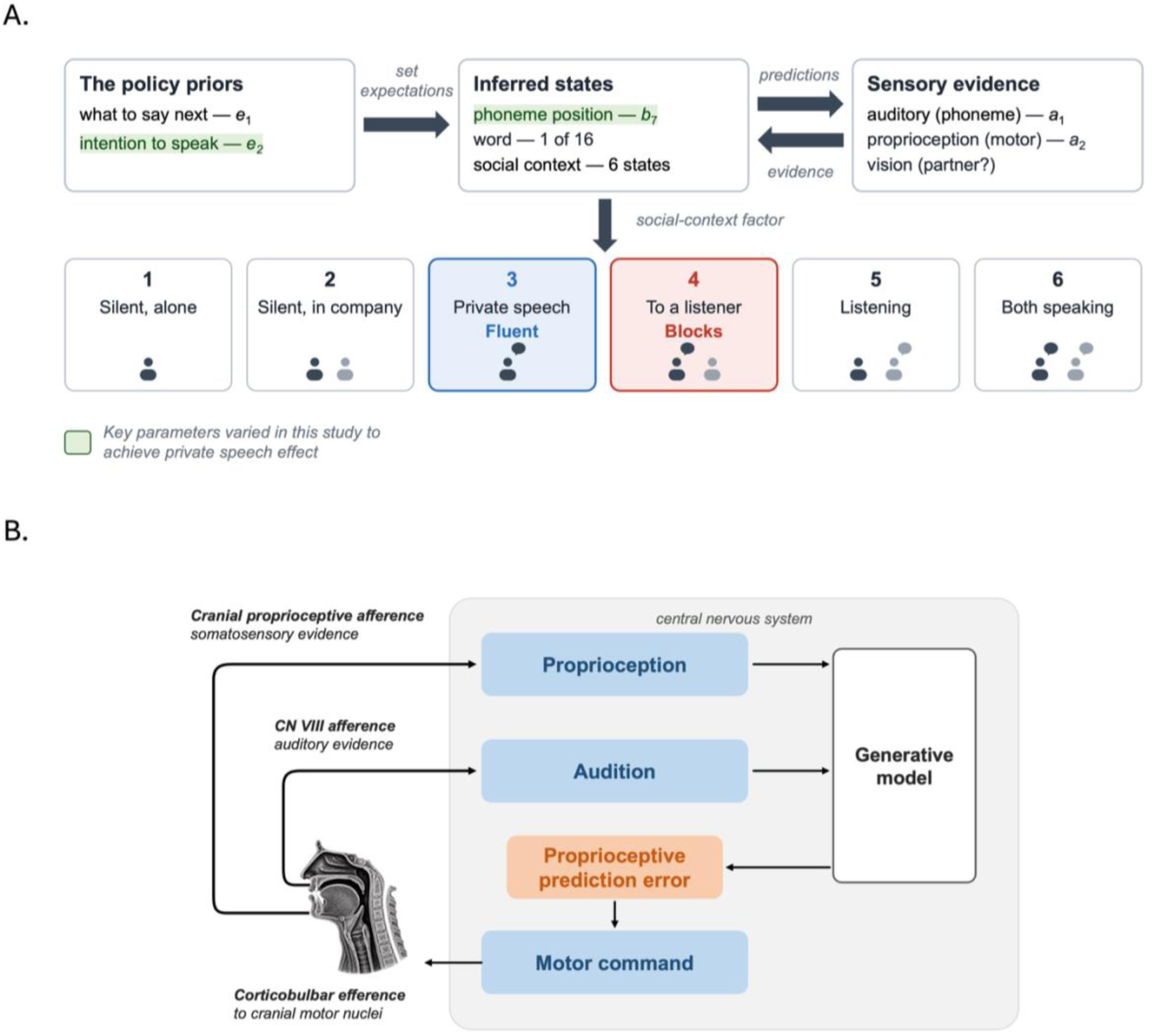
A single active-inference model links social, sensory, and motor states. **(A)** Policy priors over what to say next (*e*₁) and intention to speak (*e*₂) set expectations that guide inference over three hidden state factors: the six-state social context, the current word from a vocabulary of 16, and the current position in its phoneme sequence. The precision of transitions through the phoneme sequence is controlled by *b*_7_. These states send top-down predictions to four sensory modalities: auditory (*a*₁), two proprioceptive channels (*a*₂), and vision (partner present), and receive bottom-up evidence in return. Speech is generated when proprioceptive predictions drive motor action. The bottom row shows the six states of the social context factor. The two conditions differ only in where the agent starts, in private speech (state 3), which remains more fluent, or in speech to a listener (state 4), which produces more stuttering-like events. Green shading identifies *e*_2_ and *b*_7_, the regime-defining parameters varied in the parameter sweeps; blue and red shading identifies the condition-specific initial states. *e*₁ was set below its reference value to create graded word-transition probabilities, providing the variation in surprisal required for the information-value analysis. Parameter definitions and values are provided in Table 1. **(B)** Descending predictions from the generative model are compared with incoming proprioceptive signals, and the resulting prediction error is resolved by motor commands that move the vocal tract towards the predicted configuration. The consequences of that movement return as proprioceptive and auditory evidence, which update the generative model. Arrows to and from the generative model correspond to the prediction and evidence arrows in (A). The panel shows the sensorimotor loop for self-produced speech; visual evidence about the partner enters the generative model as shown in (A).

**Table 1:** Model parameters.

| Parameter | Interpretation | Reference value for<br>fluent speech<br>(Parr et al., 2026) | Value for stuttering-like event |
| --- | --- | --- | --- |
| $e_1$ | Inverse temperature for policy selection.<br>Controls how decisively the agent selects what to say next. | 8 | 0.94 |
| $e_2$ | Prior probability of choosing to speak. Values near 0.5 indicate near-equipose between speaking and silence. | 1.0 | 0.56 |
| $b_7$ | Precision of orbit state transitions.<br>Controls the reliability of internal timing for phoneme sequencing. | 1.0 | 0.95 |
| $b_1$ | Probability that another person arrives | 0.10 | 0.10 |
| $b_2$ | Probability that another person leaves | 0.10 | 0.10 |
| $b_3$ | Probability that, if present and agent is silent, the other person starts speaking | 0.80 | 0.99 |
| $b_4$ | Probability that the other person stops speaking while agent is silent | 0.40 | 0.40 |
| $b_5$ | Probability that, if present and agent is speaking, the other person starts speaking (interruption) | 0.40 | 0.01 |
| $b_6$ | Probability that the other person stops speaking while agent is speaking | 0.80 | 0.80 |
| $a_1$ | Precision of auditory likelihood mapping | 1.0 | 1.0 |
| $a_2$ | Precision of proprioceptive likelihood mapping | 1.0 | 1.0 |

The agent perceives the world through four sensory modalities: auditory (which phoneme is being heard), two proprioceptive channels (pharyngeal configuration and airflow), and visual (whether a conversational partner is present). It controls two action dimensions: one for selecting which path to follow through word-space (i.e., what to say next), and another for choosing whether to initiate or cease speaking. Action selection combines a prior (i.e., context-insensitive) over policies with the expected free energy of each policy, which scores how well that policy is expected to resolve uncertainty and satisfy the agent’s preferences. Only the epistemic component is relevant to our application, as we assume no preferences between alternative sensory inputs, so policies are evaluated purely by how much uncertainty they are expected to resolve. Motor output arises as the fulfilment of proprioceptive predictions, meaning that the agent generates speech by predicting the sensory consequences of intended phonemes and acting to realise those predictions through motor control (for a full description of active inference, see Parr et al., 2022).

### 2.2. Vocabulary and word transition paths

For each simulation, a pseudo-vocabulary of 16 words was generated by randomly combining simplified English phonemes (Parr et al., 2026). The inventory comprises an articulatory grid of 25 consonants arranged by manner and place, and 12 vowels by height, frontness and length. We constrained words to a minimum of two phonemes so that every word contains at least one within-word phoneme transition. A single phoneme is simultaneously the word’s onset, its only syllable, and the whole word, so it cannot express the phoneme-sequencing dynamics (governed by *b*₇) that are central to the disfluencies we model here, and it conflates word length with syllable count. The maximum length was seven phonemes. A pseudo-grammar was generated by constructing six transition matrices, each a random permutation of a circular shift matrix, defining possible word-to-word progressions. To isolate the effect of vocabulary content from the word-transition path, we held the sequence grammar and policy priors constant across all simulations using a fixed random number generator seed (for grammar and policy prior generation), while vocabulary phoneme content was varied across 1000 different seeds. This design tested whether the private-speech effect is robust to lexical variation given a fixed underlying grammar.

## 3. Simulation setup and fluency measurement

### 3.1. Parameter specification

The model parameters were chosen to replicate the specific computational vulnerabilities hypothesised to underlie stuttering in a communicative context. The critical parameters for the private-speech effect are *e*₂, *b*₇, and the social context initial state *s*. In particular, *e*₂ controls the agent’s prior tendency to speak or remain silent, *b*₇ controls the reliability of internal phoneme sequencing, and the initial social state *s* determines whether speech begins in a private or listener-present context. A fourth parameter, *e*₁, sets how widely the agent’s prior over what to say next is spread, and it is what makes information value vary across words, thereby producing variation in information value across words. Table 1 lists all parameter values and their interpretations. The remaining parameters (*b*₁ to *b*₆) govern the dynamics of the social environment, and two of these, *b*₃ and *b*₅, play a specific role that we explain below.

### 3.2. Rationale for parameter choices

Parameter selection was a calibration exercise rather than independent validation. We identified the values of *e*₂ and *b*₇ that reproduced the private speech effect, and characterised the surrounding landscape with sweeps of both. Setting *e*₁ determines the range of information values available but not the direction of any relationship between surprisal and disfluency, so that association was not a target of fitting.

#### Prior for silence (e*₂* = 0.56)

The parameter *e*₂ encodes the agent’s prior belief about whether it should initiate speech. In the default model, *e*₂ is set to 1.0 (Parr et al., 2026), meaning the agent has full confidence that it will speak. In the stuttering regime, *e*₂ was reduced from its fluent value to 0.56. Because *e*₂ is the prior probability of choosing to speak, this represents a reduced prior tendency to initiate speech, shifting the agent from near-certain speaking towards near-equipoise between speaking and remaining silent. We do not model how this prior arises in the present model; however, one possibility, which we consider in the Discussion, is that it reflects an acquired bias, emerging through threat conditioning (LeDoux & Pine, 2016).

#### Phoneme sequencing precision (b***₇*** = 0.95)

The parameter *b*₇ controls the precision of transitions around the phonemic orbit, which is the internal clock tracking progression through the phonemes within each word. At *b*₇ = 1.0 the transition distribution is a point mass, so the agent expects to occupy a single phoneme position with full certainty, and prediction errors, the mismatch between the expected and observed sensory consequences of the current phoneme, cannot revise that belief, especially when combined with a reduced prior for initiating speech (*e*_2_). The agent stays locked on the current phoneme even though the evidence supports moving on, which can pave the path for a block. Lowering *b*₇ relaxes this rigidity, but lowering it too far may leave the agent unable to maintain a stable estimate of where it is within a word. We set *b*₇ = 0.95, which places the model on the rigidity side of the fluent window and generates a simulated stuttering-like events that broadly approximates clinically observed stuttering levels.

#### Modelling the experimental paradigm (b*₃*, b₅)

The parameters *b*₃ and *b*₅ control the agent’s beliefs about the conversational dynamics of the social environment, specifically, its expectations about whether the other person will start or stop speaking. In the previous private speech study, participants were placed in two conditions (Jackson et al., 2021). First, in the private speech condition, they were left alone and prompted to speak to themselves, for instance, describing their activity during coding to improve their working memory. During the task, participants’ speech was recorded without their knowledge. Then, in the social condition, they spoke the same words to a researcher who listened but did not interrupt. The experimenter was present and attentive, creating a communicative context, but the participant was the designated speaker. Our parameter settings aimed to replicate this paradigm in part. At *b*₃ = 0.99, the agent believes that if it falls silent while someone else is present, that person will almost certainly begin speaking. At *b*₅ = 0.01, the agent believes that if it is already speaking, the other person is very unlikely to interrupt. These settings capture this experimental paradigm in which the agent is the designated speaker in a communicative context and will not be interrupted.

#### The prior over what to say (e***₁*** = 0.94)

The parameter *e*₁ is the inverse temperature on the agent’s prior over word-transition paths, reflecting how confidently the agent commits to one continuation. The reference model sets *e*₁ = 8 (Parr et al., 2026), giving a sharply peaked prior over a single near-deterministic path. We lowered *e*₁ to 0.94 by simulation to spread the prior across several of the six paths so that continuations become uncertain and information value varies across words. Because the pseudo-words carry no lexical frequency or semantic associations, word surprisal was determined by the effective transition matrix, obtained by averaging the six path matrices weighted by the same prior the agent acts on.

### 3.3. Fluency measurement

We quantified stuttering-like events following established clinical methodology adapted for the computational output (Fig. 2A). Syllables were counted using the standard phonological convention in which each vowel phoneme constitutes one syllable, with a floor of one syllable for any word containing no vowel. For each word in the 16-word pseudo-vocabulary, the number of vowel phonemes was counted, and the total for each simulation was summed over the words the agent intended, taken from its posterior over the word factor at the onset of each audible stretch, so that a word truncated by a block still contributes its full syllable count.

**Figure 2A:**
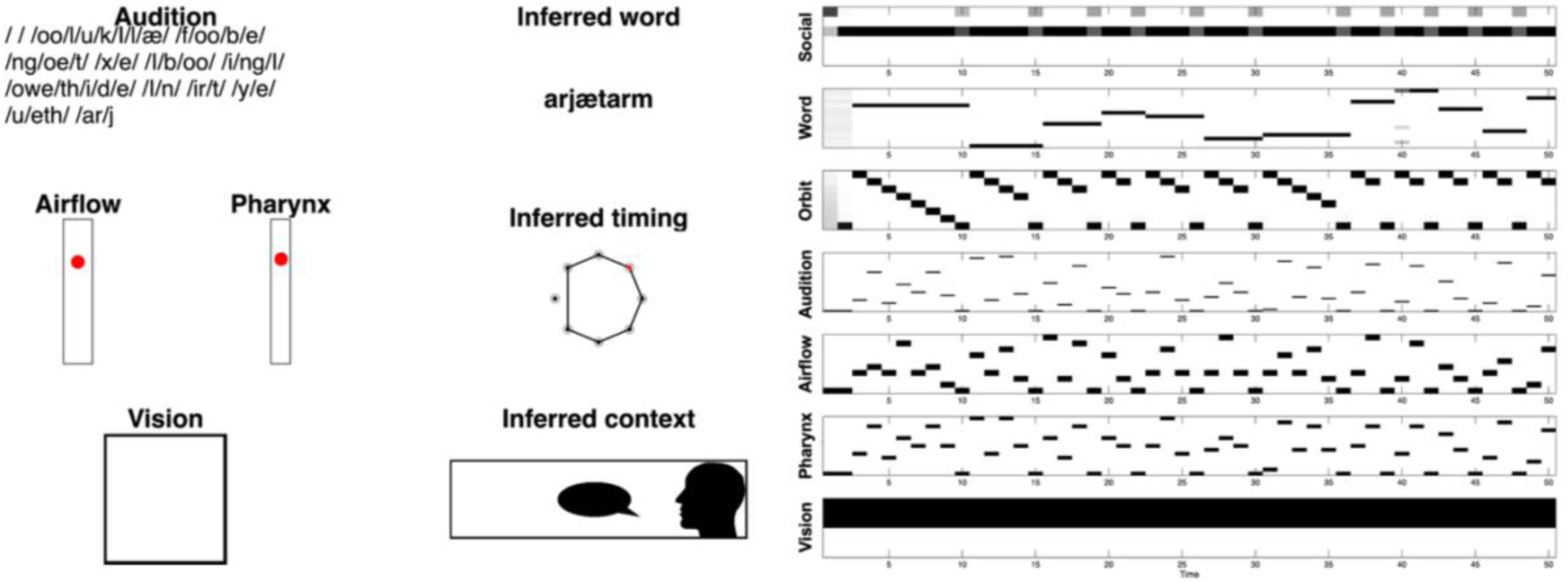
Fluent private speech. The left panel shows the accumulated auditory output as a continuous stream of phonemes enclosed in slashes (/oo/l/u/k/I/l/æ/ /f/oo/b/e/ /ng/oe/t/ …), with no prolonged gaps. Twelve words were produced, separated by single silent timesteps. Airflow and pharynx show active proprioceptive states. There is no agent in the vision box since there is no communication partner, and the inferred context box correctly shows the single agent speaking on their own. The right panel shows belief heatmaps over 50 timesteps. Black represents probability 1, white probability 0, and grey intermediate probabilities. The Word row shows crisp transitions between successive words. The orbit states align with the starts of each word. The Audition row shows dense, uninterrupted phoneme observations compatible with the corresponding proprioceptive outcomes. See Parr et al., (2026) for a detailed explanation of these plots.

#### Blocks

Silence before the first phoneme of a simulation was not scored as a block (Fig. 2B). Although blocks can occur at utterance onset in real speech, the model’s initial silence cannot be distinguished from the time the simulation takes to settle, so scoring it would conflate the two. This silence appears to be a feature of how the simulation begins rather than a product of the stuttering mechanism, since onset latency was comparable in the two conditions (2.25 and 2.29 timesteps in social and private speech) despite a more than sevenfold difference in the number of stuttering-like events.

**Figure 2B:**
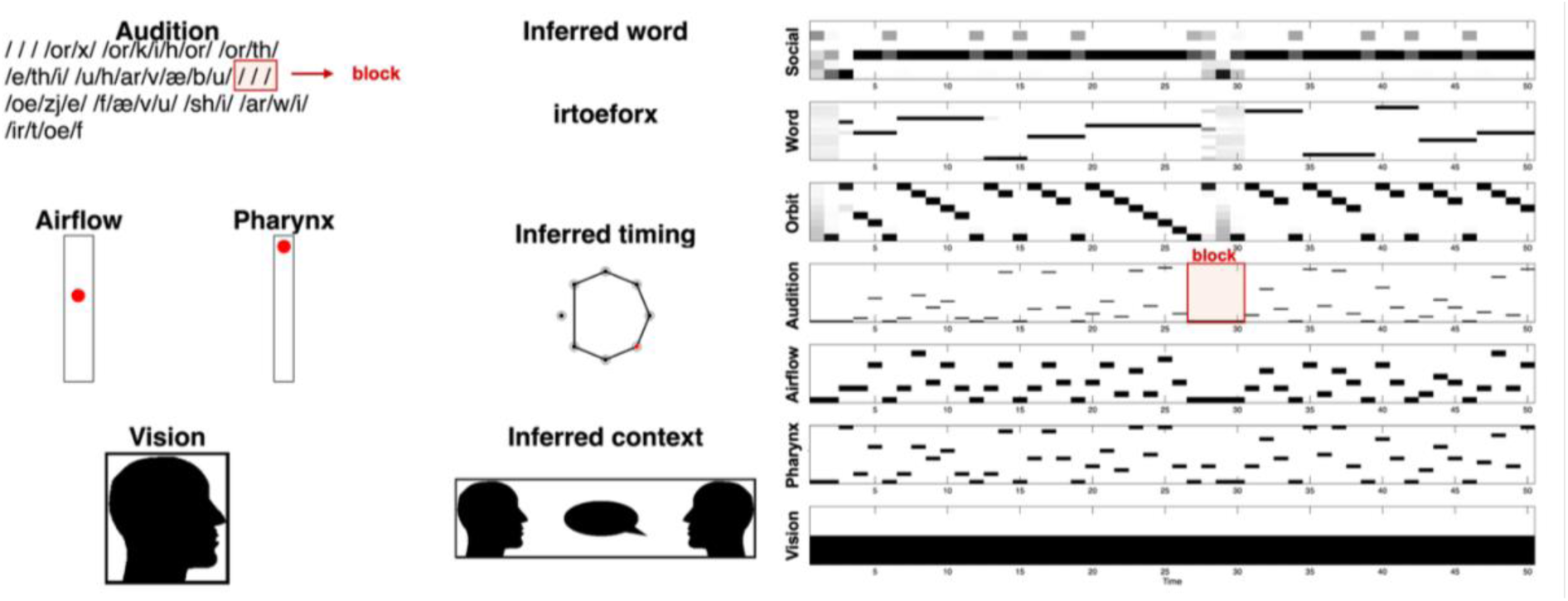
Blocks during social speech. The auditory output shows a gap in the phoneme stream where the agent produces silence, here spanning four timesteps between the fifth and sixth words. In the right panel, the Audition row shows blank columns lasting 2 or more timesteps, which mark the block. The Word row loses its sharp transitions across the same interval as the belief about which word is being produced becomes uncertain, and the Orbit row halts its progression before resuming. The Vision row is black since a communication partner is present.

#### Repetitions

Consecutive identical utterances (part-word and whole-word repetitions) were counted as a single stuttering-like event per cluster, consistent with clinical practice where a sequence of “/e/zj/ /e/zj/” counts as one instance of repetition regardless of the number of iterations (Fig. 2c). The model’s discrete phoneme output does not distinguish a sustained phoneme from a re-initiated one, so prolongations were not classified.

**Figure 2C:**
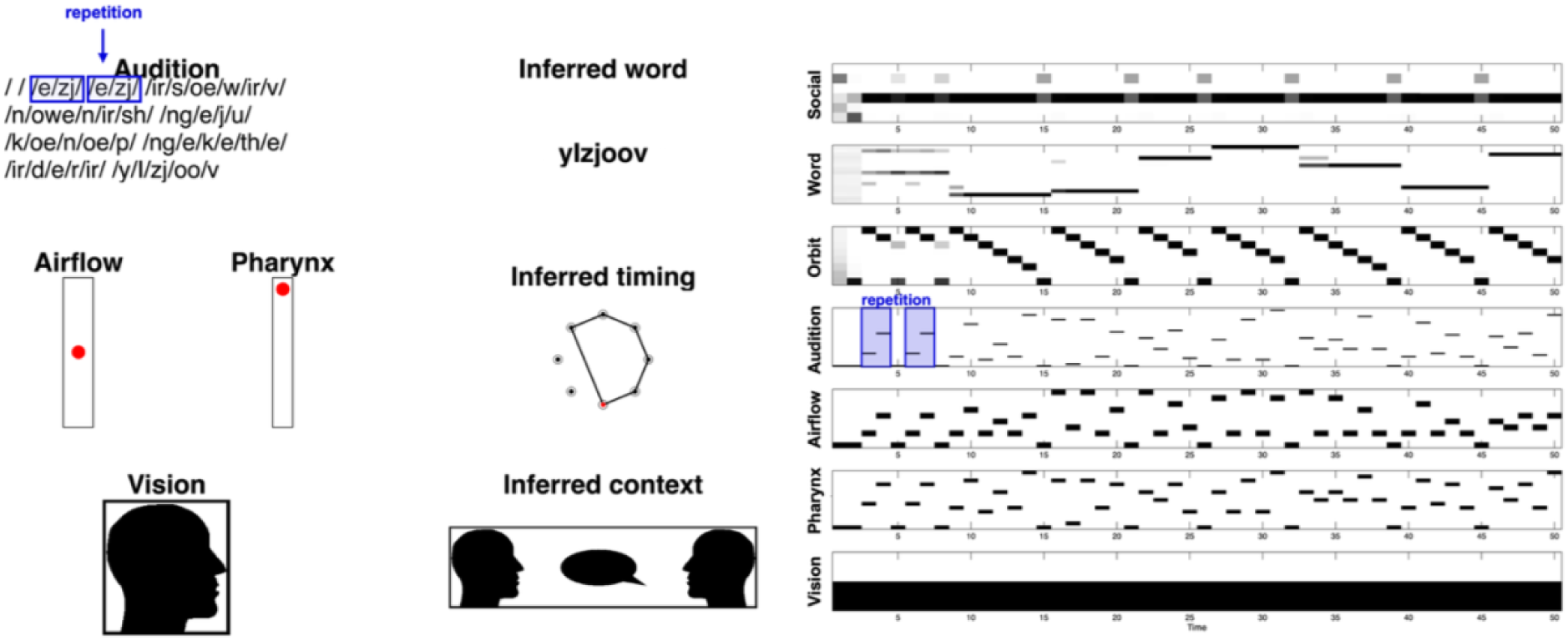
Repetition during social speech. The auditory string shows the same phoneme sequence produced twice in succession (/e/zj/ /e/zj/), separated by a single silent timestep. In the Audition row the same pattern of observations repeats before the agent moves on to the next word, and the Orbit row runs through the same short cycle twice. This simulation contained no blocks, so the repetition is its only stuttering-like event.

#### Percent syllables stuttered (%SS)

The primary outcome measure was calculated as:

%SS = (number of stuttering-like events / total intended syllables) × 100

where stuttering-like events = blocks + repetitions, and total intended syllables were derived from the model’s beliefs about which words it intended to produce via the posterior distribution over the word factor at each timestep. Please note that we retain the clinical term %SS so that model output is comparable with reported human data, but the numerator counts stuttering-like events produced by the model. Syllable counts for each vocabulary word were determined by the number of vowel phonemes in that word. Where a word was repeated, its syllables entered the total once. For a given syllable, at most one stuttering event was counted. If both a block and a repetition were associated with the same syllable (operationalised as occurring within 3 timesteps of each other), the syllable was scored as a single stuttering event (the repetition), not two separate events.

#### Validity criteria

Simulations were classified as producing “valid speech” if they met the following two criteria: (1) at least 3 unique words, and (2) at least 3 multi-phoneme words in each condition. Simulations not meeting these criteria were classified as pathological (e.g., complete silence, single-phoneme perseveration) and excluded from statistical analysis. This is analogous to clinical practice, where disfluency measurements require a minimum sample of spoken syllables to be meaningful (Riley, 2009). Moreover, repeating the same words repeatedly may reflect a different phenomenon.

#### Simulation protocol

Each simulation ran for T = 50 timesteps. For each of the 1000 vocabulary seeds, two types of simulations were conducted:

- **Private speech** (initial state *s* = 3: speaking to oneself): no conversational partner present.
- **Social speech** (initial state *s* = 4: speaking to another): a silent conversational partner present.

### 3.4. Information value analysis

A word that is hard to predict given the preceding context carries more information, which is more “surprising” in context. In everyday conversation, this may correspond to content words and rare terms, as opposed to function words or predictable completions. For example, if one says, “Pour me a cup of hot, black,” the final word is relatively predictable, whereas the final word in “She reached into her bag and pulled out a ” is not. This means that information value can be described at two related levels: the surprisal of the specific word that actually occurs, and the uncertainty over the range of possible words that could follow next. We therefore quantified information value using surprisal and entropy.

#### Surprisal

The surprisal of a word quantifies how unexpected it was, given what came before. It is measured in bits:

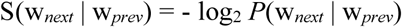

Since all lexical items in the model are pseudo-words with no real-world frequency or semantic associations, information value is defined purely by word transition probabilities. This provides a clean test of the information value hypothesis, free from lexical frequency or semantic confounds subject to individual experience. For example, if the model just produced word 5 and the probability that word 8 comes next is 0.12, then the surprisal of word 8 is -log_2_(0.12) = 3.06 bits. A highly predictable word (probability near 1) has surprisal near 0, whereas an unexpected word has high surprisal. The intended word is identified as the most probable state of the word factor at each timestep. Transition probabilities of zero produce infinite surprisal, the same issue that arises for unseen n-grams in corpus-based measures. These “off-grammar” transitions were excluded from the information value analysis, which removed 206 of 6,926 words (3.0%).

For each valid social speech simulation, we extracted the sequence of intended words from the posterior distribution over the word factor at each timestep, identified word boundaries as the timesteps where the agent’s belief about the current word changed, and computed the surprisal of each transition from the effective transition matrix. Each stuttering-like event marked exactly one word. A block marked the first word beginning after it ended, and a repetition marked the first production in the cluster. We then compared mean surprisal between the words marked stuttered and the rest. The first word of each simulation has no preceding transition and therefore no defined surprisal, so it was excluded. The analysis was restricted to social speech, since private speech produced too few stuttering-like events for a stable word-level comparison.

#### Entropy

Where surprisal tells us about a specific word, entropy tells us about the uncertainty of the whole upcoming choice. After producing a given word, how many different words could plausibly come next?

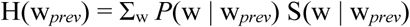

If only one word can follow (as in a fixed sequence), entropy is 0 bits. If many words are roughly equally likely, entropy is high. There are 16 words in our vocabulary; however, since the syntax is built from permuted circular-shift matrices, a word has at most six possible successors, so entropy is bounded above by log_2_(6) = 2.58 bits and not by log_2_(16) = 4. To compute these measures from the model, we needed the word-to-word transition probabilities. The model’s syntax is defined by six transition matrices (one per syntactic action), and the agent selects among these actions according to its policy prior, parameterised by a vector **E**. Consequently, we computed an effective transition matrix by averaging the six syntax matrices, weighted by the policy prior:

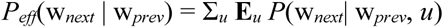

This gives a single 16 × 16 matrix where each column defines the probability distribution over possible next words, given the current word. Entropy was computed from each column of this matrix and surprisal from its individual entries, and it depends only on the preceding word. Therefore, each observed word was assigned the entropy of the column corresponding to its preceding word. As with surprisal, the first word in each simulation has no preceding transition and was excluded.

### 3.5. Word length analysis

Longer words are more likely to be stuttered in people who stutter (Howell et al., 2006; Warner et al., 2023), so we tested whether the model showed the same pattern. Word length was quantified both in phonemes (2–7 per word in the pseudo-vocabulary) and syllables (1–4). Phoneme length counts all phoneme slots, consonants and vowels, whilst syllable length counts only the vowels. For example, the 5-phoneme word “garthuh” (g-ar-th-u-h) has 2 syllables, “ar” and “u”. As above, this analysis was restricted to social speech.

### 3.6. Statistical analysis

We tested the directional hypothesis that social speech would produce higher %SS than private speech using a one-tailed paired-samples t-test, with effect sizes as Cohen’s d for paired samples. For the information value analysis, we tested whether less predictable words more likely to carry stuttering-like events. The unit of analysis was the produced word segment, and the outcome was a binary indicator of whether that segment carried a stuttering-like event. We first described the association between word-level surprisal and stuttering-like events using independent-samples t-tests and point-biserial correlations. Because the effective transition matrix was fixed across simulations, surprisal is a deterministic property of each word transition, and words are further nested within vocabularies, so the word-level observations are not independent. The inferential test was accordingly a mixed-effects logistic regression (Laplace approximation) with random intercepts for vocabulary and for transition pair, confirmed with a cluster bootstrap that resampled transitions with replacement over 5,000 iterations. To verify that the effect was consistent across vocabularies, we computed within-vocabulary point-biserial correlations for every vocabulary that varied in both measures and tested whether the mean differed from zero using a one-sample t-test on Fisher-transformed values. Eleven vocabularies with correlations of exactly ±1, where Fisher’s transformation is undefined, were excluded from that test and retained for a sign test on the direction of the correlation.

For the word length analysis we tested whether word length predicted stuttering-like events, with length measured in syllables and, separately, in phonemes. The same non-independence applies, so we again used a mixed-effects logistic regression, here with random intercepts for vocabulary and for position in the grammar. As the grammar was held fixed while the vocabulary was regenerated, each of the 16 positions carries the same transition structure in every simulation while holding a different word, so this intercept absorbs variation attaching to a position and not to a particular word. Word identity itself cannot be modelled here, since almost every combination of vocabulary and word occurs once. The grouping differs between the two models because surprisal is a property of a transition while length is a property of the word occupying a position.

A second word-length analysis was conducted at the level of the vocabulary. For each valid social speech simulation, we computed the mean and the standard deviation of word length in phonemes across its lexicon, together with the proportion of its word tokens carrying a stuttering-like event, and related these by Pearson correlations across simulations. Each simulation contributes one observation, so no clustering correction applies. We then computed the partial correlation between the standard deviation of word length and the proportion disrupted, controlling for mean word length and for total syllables produced. To test whether the same effect operates within a lexicon, each word token was assigned the absolute deviation of its phoneme length from the mean length of its own vocabulary, entered untransformed into a mixed-effects logistic regression with the same random-effects structure as above. All analyses were performed in MATLAB R2025a (MathWorks) using the Statistics and Machine Learning Toolbox. Mixed-effects logistic regressions were fitted with *fitglme* using the Laplace approximation.

## 4. Results

Among the 1,000 pseudo-vocabularies tested, 651 produced valid speech in both conditions, defined as at least three unique words, and three multi-phoneme words per simulation. The 349 excluded vocabularies failed the validity criteria predominantly in the private speech condition. Of these, 213 failed in private speech alone, 52 in social speech alone, and 84 in both (S1 Table). This may be because there is sufficient uncertainty in phoneme generation that the pair of sampled proprioceptive outcomes are incompatible with any phoneme. When the generative model assigns uniformly low probability across all candidate word sequences, no single motor plan achieves sufficient posterior confidence to be enacted. The agent resolves this uncertainty by defaulting to silence. These simulations were excluded because insufficient speech output or diversity prevents a meaningful estimate of stuttering-like event frequency. While potentially relevant for other disfluency phenomena or patterns of silence in private speech, these excluded examples are outside the scope of this analysis.

Analyses comparing private and social speech used the 651 vocabularies meeting the validity criteria in both conditions, so that each vocabulary contributed a matched pair of observations. Analyses of social speech alone, including the information value and word-length analysis, used the 864 vocabularies meeting the criteria in the social condition, since validity in the private condition has no bearing on whether social speech was scorable. Stuttering-like events occurred between words and clustered at word onset, matching the tendency for stuttering to emerge on the first sound of a word (Hubbard, 1998).

### 4.1. The model captures the private-speech effect

Across 651 valid simulations, the model generated 13,382 syllables in social speech and 13,511 syllables in private speech. Stuttering-like events occurred between words and clustered at word onset, matching the tendency for stuttering to emerge on the first sound of a word (Hubbard, 1998). Social speech generated 553 blocks and 14 repetitions, corresponding to 4.18 %SS, whereas private speech produced 75 blocks and 2 repetitions, corresponding to 0.57 %SS. Across simulations, mean %SS was significantly higher in social speech (*M* = 4.34, *SD* = 5.31) than in private speech (*M* = 0.57, *SD* = 2.27), *t*(650) = 18.19, *p* < .001, *d* = 0.71. These values broadly align with the results reported by Jackson et al. (2021), who measured stuttering rates of 0.04% of syllables in private speech and 7.46% in social speech among adults who stutter, suggesting that the model captures the private-speech effect (Fig. 3A).

**Figure 3:**
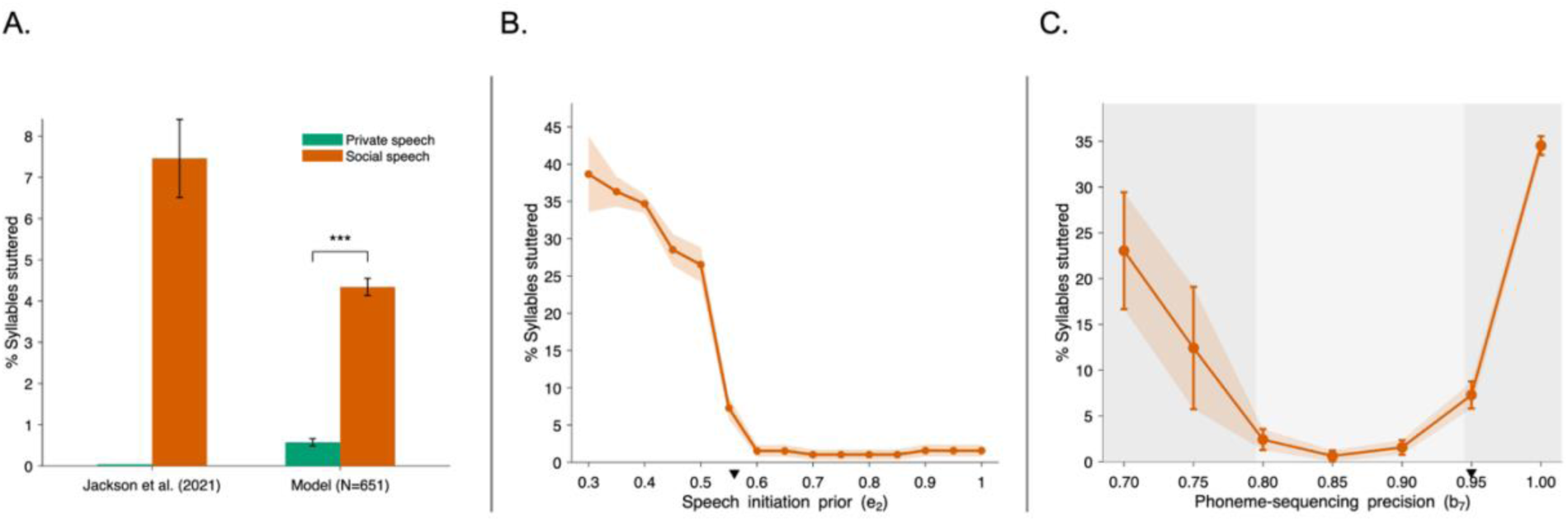
The private-speech effect and the parameter regime that produces it. **(A)** Mean percentage of syllables stuttered (%SS) in private and social speech, for the empirical data of Jackson et al. (2021) and for the model (*n* = 651 valid simulations). Jackson et al. reported 0.04% in private and 7.46% in social speech; the model produced 0.57% and 4.34%. Asterisks denote *p* < .001. For the model, %SS is the percentage of syllables carrying a stuttering-like event. **(B)** %SS in social speech as a function of the speech-initiation prior (*e*₂), swept from 0.30 to 1.00 in steps of 0.05 (ten vocabularies per value). **(C)** %SS as a function of phoneme-sequencing precision (*b*₇), swept from 0.70 to 1.00 (ten vocabularies per value). Shading marks the three regimes, unstable position tracking below 0.80, the fluency window between 0.80 and 0.95, and over-commitment above 0.95. Triangles mark the baselines (*e*₂ = 0.56, *b*₇ = 0.95). Error bars and shaded bands show standard errors.

The private-speech effect is robust even without filtering invalid simulations. Across all 1,000 vocabularies, 56.0% of social speech simulations contained at least one block or repetition compared with 11.7% of private speech simulations, and social speech averaged 1.15 such events per simulation against 0.48, *t*(999) = 10.39, *p* < .001, *d* = 0.33 (S2 Table). Applying the filter raises the same comparison to *d* = 0.78.

### 4.2. Fluency depends jointly on the initiation prior and phoneme-sequencing precision

We next examined *e*₂ as it controls the model’s prior probability of initiating speech and provides a direct test of how strongly the prior for silence contributes to stuttering-like behaviour. The sweeps use ten vocabularies per parameter value to provide a descriptive indication of the shape of each curve, whereas the main analyses use the full set of valid vocabularies. We swept *e*₂ from 0.30 to 1.00 in steps of 0.05 (social condition; Fig. 3B). All other parameters kept at baseline. Stuttering-like events followed a sharply non-linear, monotonic dose-response relationship. At the strongest silence priority setting, *e*₂ = 0.30, %SS was 38.7% (*SE* 5.1), falling to 1.0% (*SE* 0.7) by *e*₂ = 0.70. The transition was especially steep between *e*₂ = 0.50, where %SS was 26.5% (*SE* 2.3), and *e*₂ = 0.60, where it fell to 1.5% (*SE* 0.8). This range straddles the baseline, *e*₂ = 0.56 (S3 Table). Beyond *e*₂ ≈ 0.65, the curve saturated at a floor of about 1%.

We also tested the role of phoneme-sequencing precision in the model. We swept *b*₇ from 0.70 to 1.00 with the remaining parameters at their baseline values (social condition; Fig. 3C). *b*₇ produced a U-shape with an optimum operating window bordered by two different breakdowns. Below *b*₇ ≈ 0.80, the prior over phoneme positions became so diffuse that the model lost track of where it was within a word, and %SS reached 23.0% (*SE* 6.4) at *b*₇ = 0.70. Few simulations remained valid in this range, since diffuse position tracking often suppressed speech altogether, so the height of this arm is estimated less precisely than the rest of the curve (S4 Table). Above *b*₇ ≈ 0.95 the model became overly committed to its position estimate and had no flexibility to recover from momentary uncertainty, reaching 34.5% (*SE* 1.0) at the fully deterministic value of 1.00. Between these, %SS fell to 0.6% (*SE* 0.6) at *b*₇ = 0.85. There is thus a sweet spot in *b*₇ for fluent speech.

### 4.3. Social speech had equal or higher %SS in nearly all vocabularies

Across the 651 vocabularies valid in both conditions, 344 (52.8%) had higher %SS in social speech than in private speech. Private speech remained entirely fluent in 93% of the 344 vocabularies, whereas social speech contained at least one stuttering-like event. A further 297 vocabularies (45.6%) showed no difference in %SS, almost always because both conditions were entirely fluent. Only 10 vocabularies (1.5%) had higher %SS in private speech; all 10 contained stuttering-like events in both conditions. No vocabulary contained a stuttering-like event in private speech while remaining entirely fluent in social speech. Overall, private speech was entirely fluent in 91.4% of vocabularies, compared with 42.4% in social speech.

### 4.4. Words with higher information value showed more stuttering-like events

Having established that the model reproduces the communicative context aspects of stuttering, we next asked whether words of greater information value were more likely to carry stuttering-like events. Across the 864 simulations meeting the validity criteria in social speech, 6,720 words had a defined surprisal value, of which 586 (8.7%) were disrupted by a stuttering-like event and 6,134 (91.3%) were produced fluently. Disrupted words had higher surprisal (*M* = 1.46 bits, *SD* = 0.81) than fluent words (M = 1.08 bits, SD = 0.28), and the point-biserial correlation between surprisal and disruption was *r* = .29, *t*(6718) = 24.41, p < .001 (Fig. 4A). The relationship was generally graded across information values, with greater variability in the sparsely populated upper-surprisal bins (Fig. 4B).

**Figure 4:**
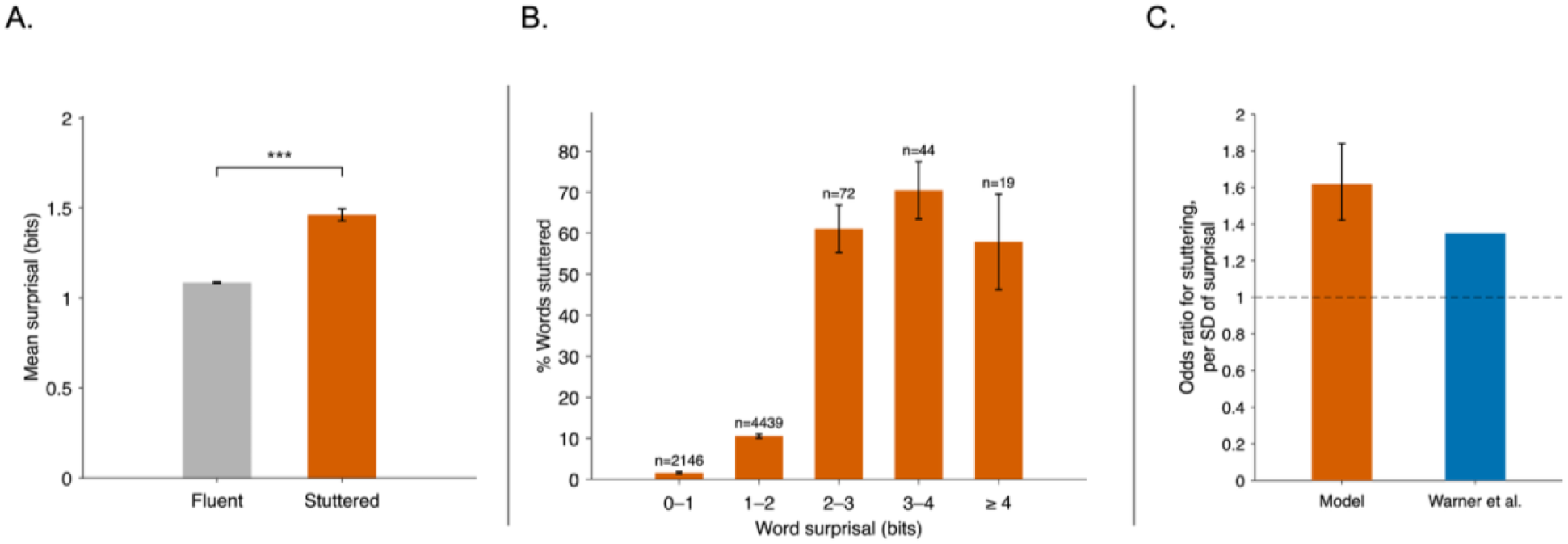
The information-value effect. Stuttered words are those carrying a stuttering-like event. **(A)** Mean surprisal (bits) for fluent (grey) and stuttered words (orange) in social speech, across 6,720 words from 864 valid simulations. Asterisks denote *p* < .001. **(B)** Percentage of words stuttered (also mentioned as disrupted in the main text) as a function of information value, defined as the surprisal (in bits) of each word given the preceding word, so that higher values mean less predictable, more informative words. Words are grouped into 1-bit surprisal bins, each spanning its lower value up to but not including the next, with the highest bin containing all words of 4 bits or more. Sample sizes are shown above each bar. Error bars show standard errors. **(C)** Odds ratio for a word carrying a stuttering-like event per one-standard-deviation increase in surprisal, estimated using a mixed-effects logistic regression with random intercepts for vocabulary and transition pair, shown alongside the corresponding estimate reported by Warner et al. (2026). The dashed line indicates the null value of OR = 1.

The grammar was held fixed across simulations, so these 6,720 words comprise only 68 distinct word transitions carrying 12 distinct surprisal values. This repetition means the words are not independent observations, so we tested the effect with methods that account for it. In the mixed-effects model, surprisal predicted stuttering with an odds ratio of 1.62 per standard deviation of surprisal, 95% CI [1.42, 1.84], *p* < .001. Fig. 4C shows this model estimate alongside the corresponding estimate reported by Warner et al. (2026). Under a word-level independence assumption, the confidence interval would have been 37% narrower on the log-odds scale. It would not have changed the conclusion. A cluster bootstrap over transition pairs, which corrects the interval for clustering without assuming a random-effects structure, placed the odds ratio between 1.47 and 2.38. Even at the low end of that range, higher surprisal raises the odds of disruption.

The effect was present within individual vocabularies as well. Of the 420 vocabularies containing both stuttered and fluent words with a range of surprisal values, 383 (91%) showed a positive correlation, *p* < .001 by sign test. Excluding eleven vocabularies with correlations of exactly ±1, the mean within- vocabulary correlation was *r* = .40, *t*(408) = 14.26, *p* < .001. This indicates that the association between surprisal and disruption is not an artefact of a few highly repeated transitions or of variation between vocabularies, and that it holds within vocabularies and survives correction for the clustered data.

The same descriptive pattern appeared in the uncertainty preceding each word. Across the same 6,720 words, entropy took eight distinct values, from 1.64 to 2.17 bits. Words disrupted by a stuttering-like event followed higher-entropy predecessors (*M* = 2.09 bits, *SD* = 0.13) than fluent words (*M* = 1.99 bits, *SD* = 0.20; pooled *r* = .14). Because entropy is fixed by the preceding word and varies only between transition-pair clusters, its estimate in a mixed-effects logistic regression with random intercepts for vocabulary and transition pair was imprecise, *OR* = 1.26 per standard-deviation increase, 95% CI [0.63, 2.53], *p* = .507. As a complementary within-vocabulary check, 376 of 423 informative vocabularies (89%) showed a positive correlation, *p* < .001 by sign test. A bootstrap resampling whole vocabularies gave a pooled *r* = .14, 95% CI [0.12, 0.16], although this interval is conditional on the fixed transition grammar. Consequently, we regard entropy as descriptively consistent with the surprisal result rather than as separate confirmatory evidence, particularly because entropy and surprisal were correlated, *r* = .48.

### 4.5. Longer words and more variable vocabularies showed more stuttering-like events

Across the 7,790 word produced by the 864 valid vocabularies, the rate of stuttering-like events rose from 8.5% for monosyllabic words to 13.7% for four-syllable words. Please note that this analysis covers more words than the information-value analysis, because a word has a length whether or not a surprisal can be computed for it, so the first word of each simulation and the off-grammar transitions are both retained here. In a mixed-effects logistic regression with random intercepts for vocabulary and for position in the pseudo-grammar, each additional syllable increased the odds of a stuttering-like event, *OR* = 1.10, 95% CI [1.01, 1.19], *p* = .033 (Fig. 5A). Phoneme count gave a comparable effect, *OR* = 1.06, 95% CI [1.01, 1.11], *p* = .010, though both effects are modest.

**Figure 5:**
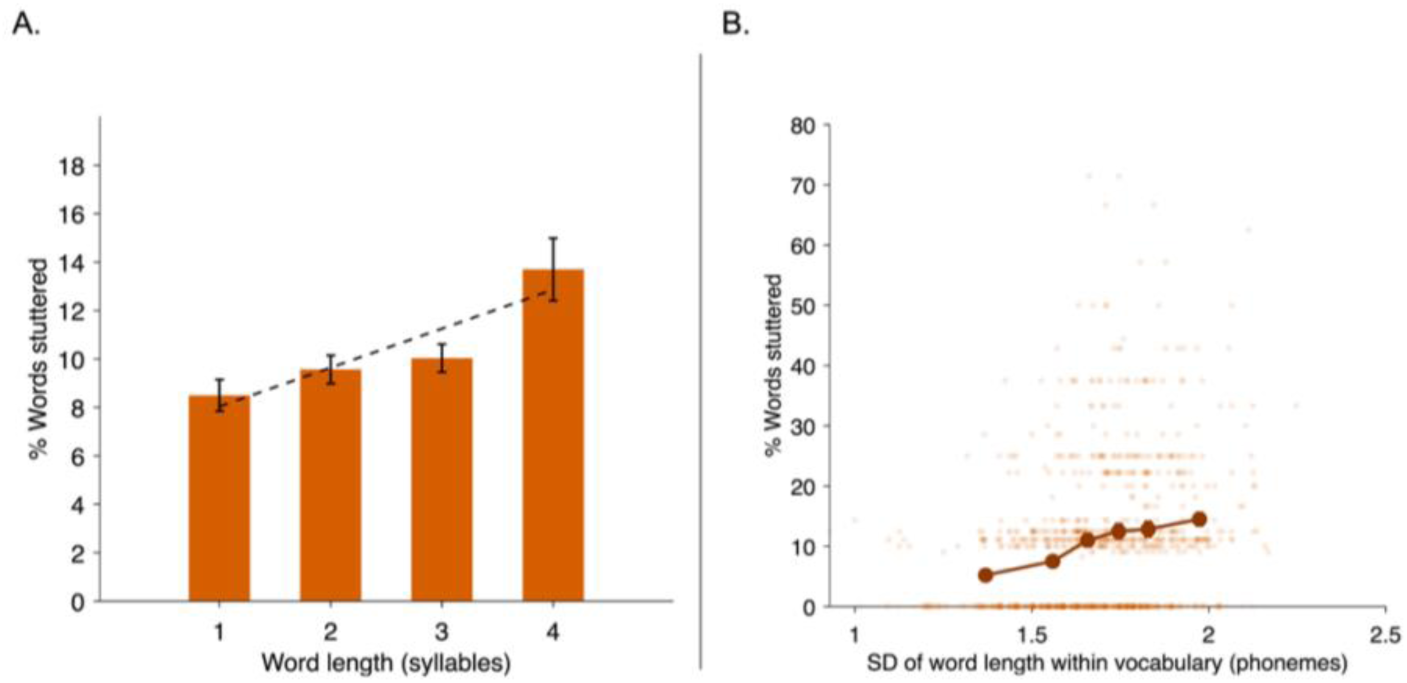
Word-length effect on stuttering-like events in simulated social speech. Both panels show the percentage of words disfluent, based on 7,790 words from 864 valid simulations in social speech. **(A)** Words grouped by syllable count, with the dashed line showing the linear trend. **(B)** Rates per vocabulary against the standard deviation of word length in phonemes within its 16-word lexicon. Faint points are single simulations, and the line shows means for six equal-sized groups. Error bars show standard errors. The percentages are discrete because a simulation produces a median of nine words, so a single stuttering-like event corresponds to roughly 11%.

A related effect was also apparent at the level of the vocabulary (Fig. 5B). Lexicons whose words varied more in length generated more stuttering-like events, with the proportion of words disfluent rising from 5.2% in the least variable vocabularies to 14.5% in the most variable, *r* = .27, *p* < .001. The relationship survived controlling for mean word length and total syllables produced, *r* = .26, and mean word length was itself a weaker predictor, *r* = .22, *p* < .001. Within vocabularies, however, words that were unusually long or short for their own lexicon were disrupted at the same rate as typical ones, OR = 1.05, 95% CI [0.96, 1.15], *p* = .29. Thus, variable vocabularies appear to be more disfluent overall, rather than containing particular vulnerable words.

## 5. Discussion

Our results show how inferred communicative context can destabilise speech initiation within an active-inference architecture. The model produced fluent private speech, greater social-speech disfluency, preferential disruption of high-surprisal words, and a word-length effect. In this synthetic agent, a prior for silence combined with rigid phoneme sequencing was sufficient to generate these effects; whether the same mechanism operates in people who stutter remains an open question.

### 5.1. A shared architecture for the private speech and information-value effects

The same architecture that generates the private-speech effect also leads to a graded sensitivity to information value. Because the agent’s prior over what to say next is spread across several word-transition paths, some words are more predictable in context than others. Less predictable words carry higher surprisal, and in the model these were more likely to produce stuttering-like events, matching the long-standing findings that less predictable words are more likely to be disfluent (Quarrington, 1965; Schlesinger et al., 1965; Soderberg, 1971; Wingate, 1979; Warner et al., 2026).

The pseudo-words used in our model had neither semantic content, nor personal history for the speaker, and information value was defined only by transition probability within the model’s word-transition structure, yet a surprisal effect emerges under these conditions. These results imply that the information value effect may not require lexical familiarity or semantic meaning. When speech is treated as an action-selection problem, word-level uncertainty becomes a plausible source of disfluency.

Both the private speech and information-value effects can be framed as the instances of the same principle. The agent withholds speech when its generative model does not fully support a confident prediction of what speaking will produce. A listener makes the consequences of resuming speech between transitions more ambiguous. Moreover, a low-probability transition makes the consequences of the next word unpredictable, since the prior over what to say next is spread. In the model, social context and lexical surprisal therefore converge on a single computational vulnerability: speech initiation becomes unstable when the consequences of speaking are difficult to predict.

### 5.2. Too much sequencing precision may be as disruptive as too little (*b*₇)

In the model, *b*₇ controls the reliability of transitions through the phoneme sequence. The sweep showed that fluency did not improve monotonically as *b*₇ increased, but followed a U-shaped pattern. At low *b*₇ the agent may not maintain a stable estimate of its position within the word. At high *b*₇, the agent can possibly become too committed to its current phoneme and less able to update when prediction error signalled that it should continue. Stuttering-like events could occur from unstable sequencing at one extreme and from excessive commitment at the other, with fluency confined to an intermediate window. Intuitively, this can be understood in much the same way as under or overfitting a model to data, and the need to obtain the right balance of accuracy while retaining a degree of tolerance to small fluctuations. Notably, *b*₇ = 1.00, the deterministic value that is the default in the reference model (Parr et al., 2026), produced the highest stuttering rate in our sweep with this parameter regime.

This high-precision regime maps onto the cortico-basal ganglia accounts in which stuttering involves impaired timing in the initiation of successive speech-motor commands (Alm, 2004; Civier et al., 2013; Chang & Guenther, 2020). *b*₇ offers a candidate computational expression of that impairment, and elevated pre-speech beta power before stuttered compared with fluent utterances (Orpella et al., 2024) is where such a correspondence would be tested.

### 5.3. Possible origins of the prior for silence (*e***₂**)

The parameter *e*₂ controls the prior probability of initiating speech, and we interpret a reduced value as a computational reflection of the implicit cost of speaking in a communicative context. This should not be read as a claim that people who stutter consciously mute themselves, nor that conscious anxiety causes speech blocks. Instead, *e*₂ might represent an implicit bias in action selection. For instance, a slight difference in the initiation system during development may produce stuttering-like repetitions, that elicit negative social evaluations of the speaker (Ferguson et al., 2019) or described by peers as being bullied or needing help (Davis et al., 2002). Constant pairing of speech with negative social evaluation could establish an implicit expectation that speaking is costly.

This might be a form of non-declarative (implicit) associative learning that operates below conscious awareness and does not require a subjective experience of fear (LeDoux & Pine, 2016). However, it may result in a shift in prior expectations. Over many years of negative social evaluation, a sense that speaking is risky and silence is safer could lead to a reduced confidence in one’s decisions to speak. Technically, this sort of prior can be developed over multiple exposures to situations in which there is ambiguity about whether to speak via the accumulation of Dirichlet counts (Fitzgerald et al., 2015). Intuitively, this can be understood as a Hebbian-like accumulation of weights that reinforce beliefs about one’s previous actions in each context. In short, repeated exposure to a situation in which one is uncertain about what to do favours development of an empirical prior that reflects this ambiguity. This interpretation aligns with claims that threat conditioning modulates stuttering (Peters & Guitar, 1991). On that account, stuttering may involve a conditioned freeze response inhibiting vocalisation under perceived threat. Whether such threat conditioning can be represented as a reduced prior for speaking is open for inquiry.

### 5.4. Length effects at the word and lexicon level

The word-length effect appeared in the same parameter regime as the private speech and information-value effects. On the EXPLAN account, disfluency arises from a mismatch between executing one word and planning the next, so that fluency fails at the junction where the plan is not ready when execution arrives (Howell & Au-Yeung, 2002). A related proposal is that competing phonological segments stay active too long during planning, leaving the speech plan error-prone (Postma & Kolk, 1993). In our model, low *b*₇ is a loose analogue, with the agent’s belief spread across adjacent positions in the sequence instead of across competing segments. Length effects do not require lexical or semantic structure, since they also appear in a nonword repetition task (Sasisekaran & Weathers, 2019), which makes a pseudo-vocabulary a reasonable place to look for one.

A further finding was that lexicons whose words varied more in length generated more stuttering-like events. Within a lexicon, however, a word’s departure from the local mean did not predict whether it was disrupted, in either direction. The effect operates at the lexicon level rather than being confined to outlier words. The agent does not know which word comes next, and the orbit’s dynamics depend on which word it is, so its expectation about when a word will start and end is a mixture over the candidates. That mixture is more coherent in terms of fluency when words are close in length and more incoherent when they vary, so the effect may be seen across the lexicon instead of on particular words. A comparable relationship has been suggested for lexical diversity, with more varied vocabulary in the language samples where a child stutters most (Wagovich & Hall, 2018), though, to our knowledge, variation in word length has not been examined.

### 5.5. Clinical and experimental implications

A potential clinical use of our model is to distinguish mechanisms that look similar in behaviour. Comparable rates of stuttering-like events occurred at a low prior for speaking and at a rigid sequencing precision, so the overt rate does not identify which setting produced it. Therefore, the model opens a route to computational subtyping, describing an individual speaker by the process beyond their speech fluency.

The model also has potential implications where fluency intervention might be more beneficial. For example, disfluency emerged when the agent inferred that a listener was present, so we can hypothesise that fluency practised without one may not transfer effectively to speech that is addressed to another agent. Training under communicative demand, of the kind a controlled social context can provide such as virtual reality exposure therapies (Freeman et al., 2018), is worth testing. For example, an external cue that brings sequencing into a fluent state under communicative context, such as speaking fluently in unison with another voice through headphones while giving a presentation (Demirel, 2025), would act on the similar parameter the *b*₇ sweeps identify. Continuous experience of fluent speech with communicative intent may eventually update the prior itself with the aim of persisting fluency, even when the cue is withdrawn.

Our second prediction concerns speech onset. In the model, blocks occur at word onsets, where the agent stays silent instead of entering a state it cannot predict. This locates the disfluency in the pre-initiation window and predicts that a neural signature of altered initiation should appear there. This is in line with neuroimaging evidence of reduced or disrupted speech-motor preparation at initiation (Mersov et al., 2016; Orpella et al., 2024). It further predicts that shifting this state immediately before speech onset may change whether a block occurs, which can be tested with phase-specific neuromodulation (Cagnan et al., 2017; Mancini et al., 2026).

Our final implication concerns experimental design. Because blocks in the model depend on an inferred listener, paradigms that leave the speaker isolated may underestimate stuttering. Eliciting stuttering in laboratory environments is known to be difficult (Jackson et al., 2020), and some neuroimaging studies have addressed this by including a listener, either through live interpersonal communication in the scanner (Toyomura et al., 2018) or through tasks built around experimenter interviews (Lu et al., 2025). The model offers one account of why such designs might help, and suggests that a communicative context is worth considering as a design variable instead of an incidental feature of the setting.

### 5.6. Limitations and future directions

The model is explanatory and has not been fitted to individual human data, so while a small set of assumptions reproduces the features of stuttering, it cannot say which combination applies to a given speaker. Fitting individual speech samples would test whether individuals of similar overt severity differ in the prior for speaking and in sequencing precision. Validating the parameters would mean using the fitted values as regressors in analyses of neuroimaging data such as pre-speech beta activity already associated with stuttering (Mersov et al., 2016; Orpella et al., 2024).

Approximately one third of the vocabularies were excluded because private speech produced too little output to score. This exclusion might not be neutral, since failure to initiate speech is close to the mechanism under study. The simulated output captured a narrower range of behaviours than clinical stuttering. The model generated mostly blocks and few repetitions, and it cannot represent prolongations, since its discrete phoneme output does not distinguish a sustained phoneme from a re-initiated one. Blocks in the model always occur at the onset of a word, which matches the word-initial bias in human stuttering (Hubbard, 1998); however, the model does not represent the minority of events that fall on mid-syllables. Future work should examine which parameter combinations produce which symptom profiles.

Finally, communicative context was treated as binary. A speaker may be alone yet imagine a listener, or record their voice knowing someone will hear it later, and stuttering still occurs under those conditions (Demirel, 2025). Extending the communicative context to distinguish a partner who is physically present, one who is imagined, and one who is present but with varied attention to speaker’s speech, would let these be tested separately.

## 6. Conclusion

The present work offers a computational account of how stuttering may emerge when speech is produced in a communicative context. Within a single parameter regime, speech was fluent when the agent inferred it was alone, became more vulnerable when it inferred a listener was present, disrupted more often on words that were harder to predict, and showed a word-length effect. By representing communicative context as an inferred state that increases unpredictability, the model reframes stuttering-like events as a possible emergent property of a socially responsive speech production system. Therefore, a unified account of stuttering must include communicative context alongside speech-motor mechanisms. Clinically, the model makes testable predictions. Within the jointly calibrated regime, different settings of the prior for speaking and phoneme-sequencing precision produced comparable levels of stuttering-like behaviour. If these parameter differences have human analogues, individuals matched on overt severity may nevertheless differ mechanistically, which could predict differential responses to intervention.

## Code availability

Simulation code and analysis scripts are available at https://github.com/Birtan-Demirel/Stuttering-Active-Inference. The two main-condition scripts each run 1,000 pseudo-vocabularies over 50 timesteps using the parameter values in Table 1. The parameter-sweep scripts run ten vocabularies at each tested different values of *e*₂ or *b*₇. All scripts detect blocks and repetitions, calculate %SS, and apply the validity criteria described above. The generative model follows Parr et al. (2026) and the reference implementation at https://github.com/tejparr/Computational-Neurology.

## Supporting information

**S1 Table.** Classification of all 1,000 simulated vocabularies. Outcome of every simulation in each condition, classified as valid speech, silence, perseveration or insufficient diversity.

**S2 Table.** Sensitivity of the private-speech effect to the validity filter. The private speech comparison computed with and without the validity filter, using stuttering-like event counts and the percentage of syllables stuttered.

**S3 Table.** Dose-response analysis for *e*₂, the prior probability of initiating speech. Mean percentage of syllables stuttered at each of 15 values of *e*₂ in the social condition, with standard errors and the number of simulations meeting the validity criteria.

**S4 Table**. Dose-response analysis for *b*₇, the precision of phoneme sequencing. Mean percentage of syllables stuttered at each of 19 values of *b*₇ in the social condition, with standard errors and the number of simulations meeting the validity criteria.

