## Supplemental Data 1 for "Why Speech Motor Blocks Emerge in a Communicative Context: An Active Inference Model of Stuttering"

### S1 Table: Classification of all 1,000 simulated vocabularies.

| Outcome category | Private speech | Social speech |
|---|---|---|
| Valid speech | 703 | 864 |
| Silence | 257 | 86 |
| Perseveration | 40 | 50 |
| Insufficient diversity | 0 | 0 |
| Total | 1,000 | 1,000 |

Categories are mutually exclusive and were applied to each condition separately. Valid speech, at least three unique utterances and at least three multi-phoneme utterances; Silence, fewer than three utterances produced; Perseveration, at least three utterances but fewer than three unique utterances; Insufficient diversity, at least three unique utterances but fewer than three multi-phoneme utterances. Analyses comparing the two conditions used only vocabularies meeting the criteria in both, of which there were 651. Among the 349 excluded vocabularies, 213 failed in the private condition only, 52 in the social condition only, and 84 in both.

### S2 Table: Sensitivity of the private-speech effect to the validity filter.

| Analysis | *n* | Social %SS | Private %SS | *t* | *p* | *d* |
|---|---|---|---|---|---|---|
| Stuttering-like events, all vocabularies | 1,000 | 1.15 (1.96) | 0.48 (1.98) | 10.39 | < .001 | 0.33 |
| Stuttering-like events, filtered | 651 | 0.87 (0.96) | 0.12 (0.47) | 19.81 | < .001 | 0.78 |
| %SS, filtered (main analysis) | 651 | 4.34 (5.31) | 0.57 (2.27) | 18.19 | < .001 | 0.71 |

Values are means across vocabularies with standard deviations in parentheses. Stuttering-like events are the sum of blocks and repetitions in a simulation, the only fluency measure defined for every run, since vocabularies producing no speech have no syllable denominator. Tests are one-tailed paired-samples t-tests. *d* = Cohen's d for paired samples. The unfiltered comparison is conservative. Excluded vocabularies are divided into two separated paths: runs that produce almost nothing, with a median of zero syllables in each condition, and perseverative runs that loop on a single phoneme separated by silences, each silence scoring as a block. The 27 perseverative runs in private speech carry 64% of its unfiltered event total. Social speech produced more stuttering-like events whether or not the filter was applied.

#### S3 Table: Dose-response analysis for $e_2$, the prior probability of initiating speech.

| $e_2$ | Mean %SS | SE | Valid simulations |
|---|---|---|---|
| 0.30 | 38.66 | 5.09 | 8 |
| 0.35 | 36.31 | 2.01 | 8 |
| 0.40 | 34.66 | 1.24 | 6 |
| 0.45 | 28.52 | 2.14 | 10 |
| 0.50 | 26.49 | 2.32 | 10 |
| 0.55 | 7.28 | 1.47 | 10 |
| 0.60 | 1.53 | 0.78 | 10 |
| 0.65 | 1.53 | 0.78 | 10 |
| 0.70 | 1.03 | 0.69 | 10 |
| 0.75 | 1.03 | 0.69 | 10 |
| 0.80 | 1.03 | 0.69 | 10 |
| 0.85 | 1.03 | 0.69 | 10 |
| 0.90 | 1.59 | 0.81 | 10 |
| 0.95 | 1.56 | 0.80 | 10 |
| 1.00 | 1.56 | 0.80 | 10 |

Ten vocabularies were simulated at each of 15 values of $e_2$ in the social condition. All other parameters were held at their baseline values, $e_1 = 0.94$, $b_7 = 0.95$, T = 50. Values are means across the simulations meeting the validity criteria at each value, with the standard error of the mean. Fewer than ten simulations were valid at $e_2 = 0.30$, 0.35 and 0.40, where a strong prior for silence frequently suppressed speech entirely. Identical means at adjacent values reflect a floor in stuttering above $e_2 \approx 0.60$, beyond which further increases in the prior for speaking produced no additional change. The baseline used throughout the paper, $e_2 = 0.56$, lies between the two steepest points of the curve and was not itself part of the sweep. SE = standard error

#### S4 Table: Dose-response analysis for $b_7$, the precision of phoneme sequencing.

| $b_7$ | Mean %SS | SE | Valid simulations |
|---|---|---|---|
| 0.70 | 23.04 | 6.37 | 2 |
| 0.72 | 24.93 | 5.21 | 6 |
| 0.74 | 12.47 | 6.67 | 6 |
| 0.75 | 12.42 | 6.68 | 6 |
| 0.76 | 13.79 | 6.87 | 6 |
| 0.78 | 11.45 | 5.91 | 8 |

| | | | |
|---|---|---|---|
| 0.80 | 2.41 | 1.15 | 7 |
| 0.82 | 1.47 | 0.95 | 7 |
| 0.84 | 0.66 | 0.66 | 8 |
| 0.85 | 0.62 | 0.62 | 8 |
| 0.86 | 0.53 | 0.53 | 9 |
| 0.88 | 1.56 | 0.80 | 10 |
| 0.90 | 1.56 | 0.80 | 10 |
| 0.92 | 1.59 | 0.81 | 10 |
| 0.94 | 1.53 | 0.78 | 10 |
| 0.95 | 7.28 | 1.47 | 10 |
| 0.96 | 7.31 | 1.43 | 10 |
| 0.98 | 29.32 | 2.11 | 10 |
| 1.00 | 34.52 | 1.04 | 9 |

Ten vocabularies were simulated at each value of $b_7$ in the social condition, with all other parameters held at their baseline values, $e_1 = 0.94$, $e_2 = 0.56$, T = 50. Values are means across the simulations meeting the validity criteria at each value, with the standard error of the mean. The value 0.95 is the baseline used throughout the paper. Validity dropped as low as two simulations below $b_7 = 0.80$, where diffuse position tracking often suppressed speech altogether, so the left arm of the curve is estimated less precisely than the right. Stuttering was lowest between $b_7 = 0.84$ and 0.86 and rose steeply on both sides, reaching 34.52% at the fully deterministic value of 1.00. SE = standard error.
